# ATP Mediates Yeast Cell-Cell Communication

**DOI:** 10.1101/2020.07.02.185231

**Authors:** Michael Caplow

**Affiliations:** Department of Biochemistry and Biophysics, University of North Carolina, Chapel Hill, NC 27599

## Abstract

Yeast secrete ATP in response to glucose, a property with previously unknown functional consequence. In this report, we show that extracellular ATP is a signal for growth of surrounding cells. The ATP signaling behavior was serendipitously uncovered by finding reduced toxicity of an inducible, dominant-lethal form of alpha tubulin (tub1-828); lethality was dramatically reduced in cultures grown at high, compared to low cell density. Reduced cell death at high cell density occurred because the mutant tubulin’s deleterious effect of mitosis was reduced such that the rate of chromosome loss/cell division was lower (18-fold) in cultures inoculated with a high density (75,000/ml) compared to a low density (1,000/ml) of cells. The sparing effect of growth at high cell density could be replicated by co-culturing a low number of cells (3440/5 ml) that expressed tub1-828, with a high number of cells (2.3 E6/5 ml) that did not express the mutant protein. In this condition toxicity was reduced at high cell density apparently because it produced a sufficient concentration of a secreted growth substance, such that the mutant protein was rapidly diluted by synthesis of wild-type alpha tubulin (TUB1). Enhanced growth at high cell density was confirmed by fluorescence-activated cell sorting (FACS) analysis after DNA staining, which showed that the rate of the G1-G2 transition was faster for cells at high density. ATP replaced the need for high cell density for resistance to tub1-828, and stimulated the transition from G1 to G2 in cells at low density. This newly discovered quorum sensing response in yeast, mediated by ATP, indicates that yeast decision-making is not entirely autonomous.

## Introduction

In multicellular organisms, extracellular ATP functions as a signaling molecule for stimulation of cell growth and regulation of development. Signaling occurs by the binding of ATP to specific purinergic receptors. The unicellular yeast *Saccharomyces cerevisiae* also secretes ATP by way of a specific vesicular pathway ATP (Boyum and Guidotti, 1997; Esther et al., 2008, Peters et al., 2016). However, *S. cerevisiae* apparently lacks any homolog of the known purinergic receptors; thus, the function of yeast ATP efflux is unknown. In this report, we present evidence that extracellular ATP is a signaling molecule that modulates yeast cell growth.

Evidence for extracellular ATP signaling in *S. cerevisiae* came from the cell growth characteristics of a tubulin mutant (Anders and Botstein 2001; tub1-828) in studies expected to elucidate further the mechanism for GTP hydrolysis required for microtubule assembly and dynamics (Caplow et al., 1985; Caplow and Reid 1985; Caplow and Shanks J, 1996). The tub1-828 mutation, which is in the GTPase activating domain of the alpha subunit, has a dominant lethal phenotype that results from formation of abnormally stable microtubules that cause cell death. Unexpectedly, we found that the lethality of tub1-828 was dramatically reduced when cells were inoculated into medium at a high cell density. This density-dependent rescue of tub1-828 suggested that signaling by an extracellular substance had stimulated expression of the wild type TUB1 gene present in the same strain, and overexpression of wild type TUB1 diluted the assembly of tub1-828 into microtubules. It had been established that synthesis of wild-type TUB1 can suppress tub1-828 toxicity (Anders and Botstein 2001, Fig.1). We surmised that a sufficiently high concentration of ATP from the high density of cells was the extracellular substance that modulated cell growth and tub1-828 toxicity. We confirmed this conjecture by observing enhanced growth of cells expressing tub1-828 plated on solid media containing ATP. These results and those from related studies provide evidence that ATP secreted by yeast generate a growth signal for surrounding cells.

**Fig. 1.**
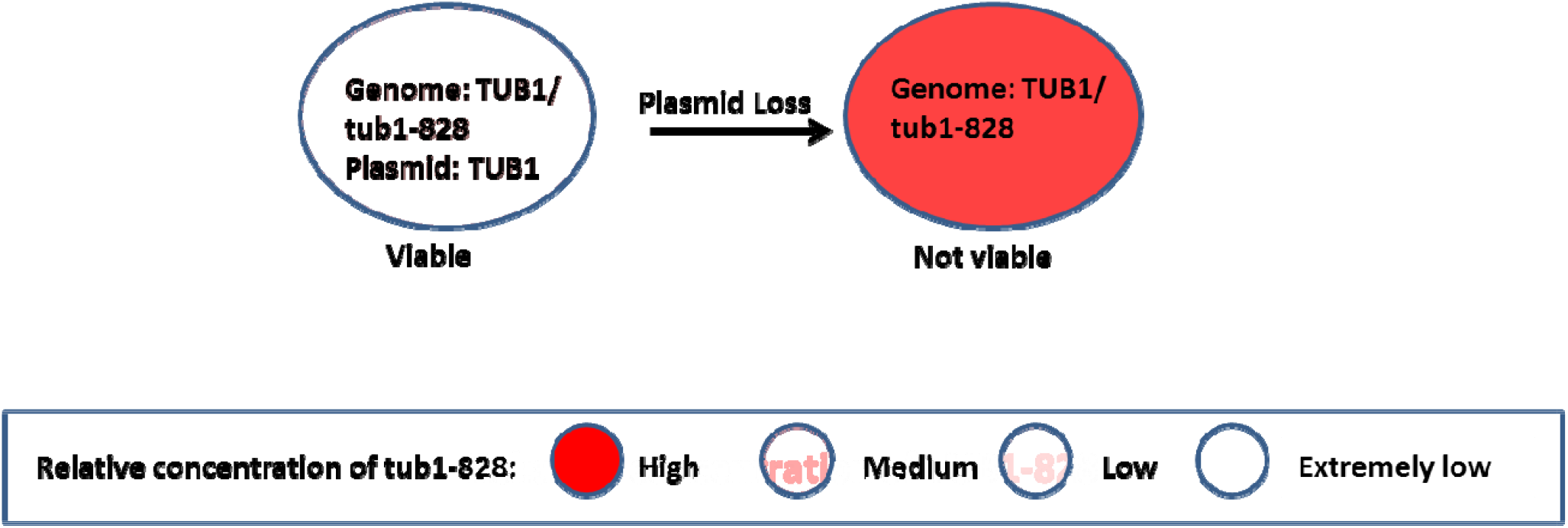
tub1-828 is only toxic at high concentration. DBY6596, a diploid heterozygote with TUB1 and tub1-828 is viable as long as it contains pRB326, a CEN plasmid containing TUB1 expressed from a native promoter.

### Experimental Methods

#### Insertion of tub1-828 into the yeast genome

To produce a plasmid for integration of tub1-828 into the yeast genome, the following modifications were made to the episomal plasmid pCM190 (obtained from Euroscarf). First, the tet-off promoter of pCM190 (5) was replaced with a tet-on promoter, contained in a BamHI/EcoRI fragment from plasmid pCM252 (from Euroscarf). Second, the tub1-828 coding sequence (from DBY2963, obtained from Dr. David Botstein) was inserted at the BamHI site to place it under the control of the tet-on promoter.

Third, after a cytosine residue that flanks and prevents cleavage at the Cla1 site in pCM190 had been deleted, the pCM190 plasmid containing tub1-828 was PCR-amplified so as to delete the 2-micron element that confers episomal replication. Fourth, the 2 micron-less plasmid was used as a template for PCR with a primer containing 50 bp of 5’-untranslated TRP1 sequence and a reverse primer contained 50 bp immediately following the TRP1 stop codon; the resultant linear PCR product containing the tet-on tub1-828 gene was used to transform strain DBY3363 cells that had been modified as follows.

DBY3363 is a haploid yeast strain that contains a multiply marked supernumerary copy of Chromosome III. The extra chromosome that enables chromosome loss to be observed easily on various media. Two modifications to DBY3363 were made before introducing into its genome the tet-on tub1-828 PCR product (see above) into its genome. First, we disrupted the ura-3-52 allelle in DBY3363 with a hygromycin resistance gene (from pAG2). This disruption prevented repair of ura3-52 and enabled selection for integration of the URA3-containing tet-on tub1-828 pCM190 plasmid. Second, the gene encoding the reverse tetracycline-controlled transcription activator (rtTa), which activates the tetracycline response elements only in the presence of doxycycline was introduced into DBY3363. The rtTA gene was present in plasmid pCM242; this plasmid was modified to contain the yeast LYS2 gene and the LYS2 gene of DBY3363 was disrupted by insertion of nourseothricin resistance gene, thereby providing a means to select for genomic insertion of the LYS2-containing pCM242 plasmid. Finally, the tet-on tub1-828 construct was amplified by PCR with primers containing flanking sequences of TRP (see earlier) and the PCR product was used to replace the TRP1 gene of DBY3363, selecting for URA^+^ cells.

The nucleoside diphosphate kinase gene YNK1 was inactivated by insertion of TRP1.

### Analysis of Cell Behavior

Stationary-state cells were generated by growth of a large inoculum for 24-72 hours in YPD liquid media in the absence of doxycycline; log-phase cells were obtained from overnight growth of a small inoculum. For growth studies the inoculum was quickly washed with media into which they were to be grown. Induction of tub1-828 was in liquid or on agar plates with doxycycline (5 ug/ml), and the extent of growth in liquid media was determined with a hemocytometer. DNA in cells was stained as described (Haas and Reed, *2002*), and images were collected with a BD LSR II Flow Cytometer. FACS analysis was performed in the absence of tetracycline, so that formation of tub1-828 did not influence the results.

The effect of tub1-828 expression was determined in liquid YPD-tetracycline or on agar plates with tetracycline and, in some cases, with nucleotide; these substances were added after autoclaving and they did not alter the pH of the media. Comparisons of cell growth were performed with plates prepared on the same day and stored under identical conditions. The number of cell divisions after growth on agar was estimated from measurement of colony diameter using the reticle in a low-power microscope, assuming that the colony and individual cells (5 μm diameter) were spherical, with cells packed so that they occupied 71% of the volume; this value is between the 63% for random close packing and 74% for hexagonal close packing. Stimulation of growth was quantitated from a histogram, using the number of cells that had divided n-times to calculate the number of cells that would be contained in 100 colonies. Evidence that this method was accurate was finding a similar doubling time when growth was measured with log-phase DBY3363 cells plated on YPD-agar (1.77 h doubling time) and when a hemocytometer was used to analyze the same cells grown in liquid YPD (1.55 h doubling time).

ATP secretion was measured with a LB953 AutoLumat luminometer, using the ApoSENSOR ATP Cell Viability Kit for analysis of samples after removing cells by centrifugation.

### Materials

Cryptopleurine was a generous gift from Dr. Paul Savage at CSIRO and was used as described (Hoyt et al., 1990).

## Results

### Cells for Studies of Growth Effects

<u>Because tub1-828 expression results in plasmid loss (unpublished results), studies of its effect on cell growth were with DBY3363 cells that had been modified to express tub1-828 from a tet-on promoter</u> (Belli et al., 1998) inserted in the TRP1 sequence in Chromosome IV. This haploid strain contains a multiply marked supernumerary copy of Chromosome III, and cells that lose one of the extra chromosomes form red colonies that are resistant to cryptopleurine (Hoyt et al., 1990); cells that lose any other chromosome are not viable. This property allowed determination of the effect of tub1-828 on chromosome loss.

### Effect of tub1-828 expression

tub1-828 toxicity was determined with cells grown in the presence of tetracycline on solid and in liquid media. Most cells grown on agar could undergo 2-12 divisions in 48 hours, despite the fact that tub1-828 had been identified as conferring a dominant-lethal phenotype. Cells that divided 2-12 times produced microscopic colonies; however, a highly variable fraction of cells divided about 20 times and formed 1 mm diameter macrocolonies (Fig. 2A,B).

**Fig. 2A.**
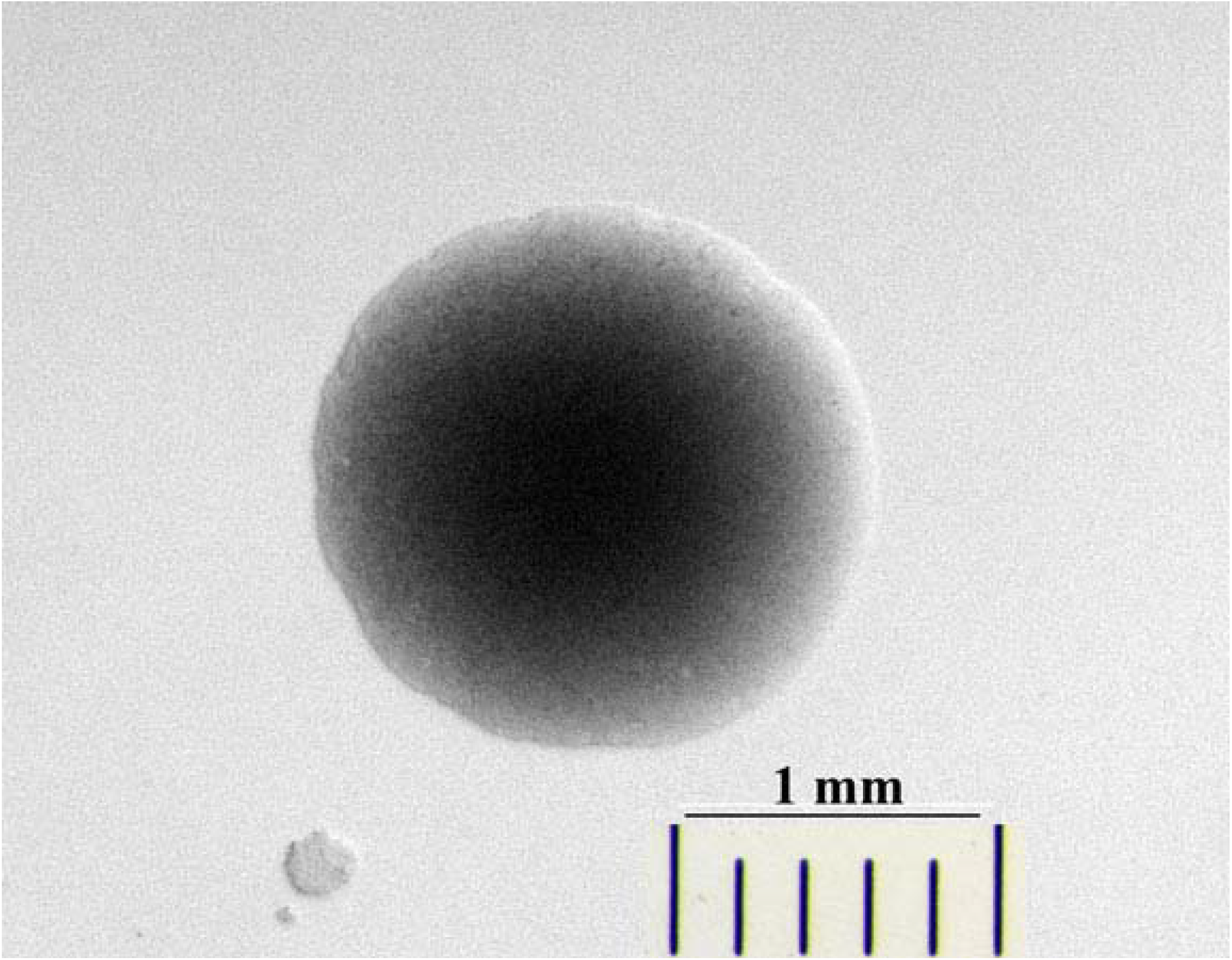
Formation of micro and macrocolonies with cells (MCY046) expressing tub1-828. Colonies with a 1 mm diameter colonies contain about one million cells. Note the variation in the size of the three colonies.

**Fig. 2B.**
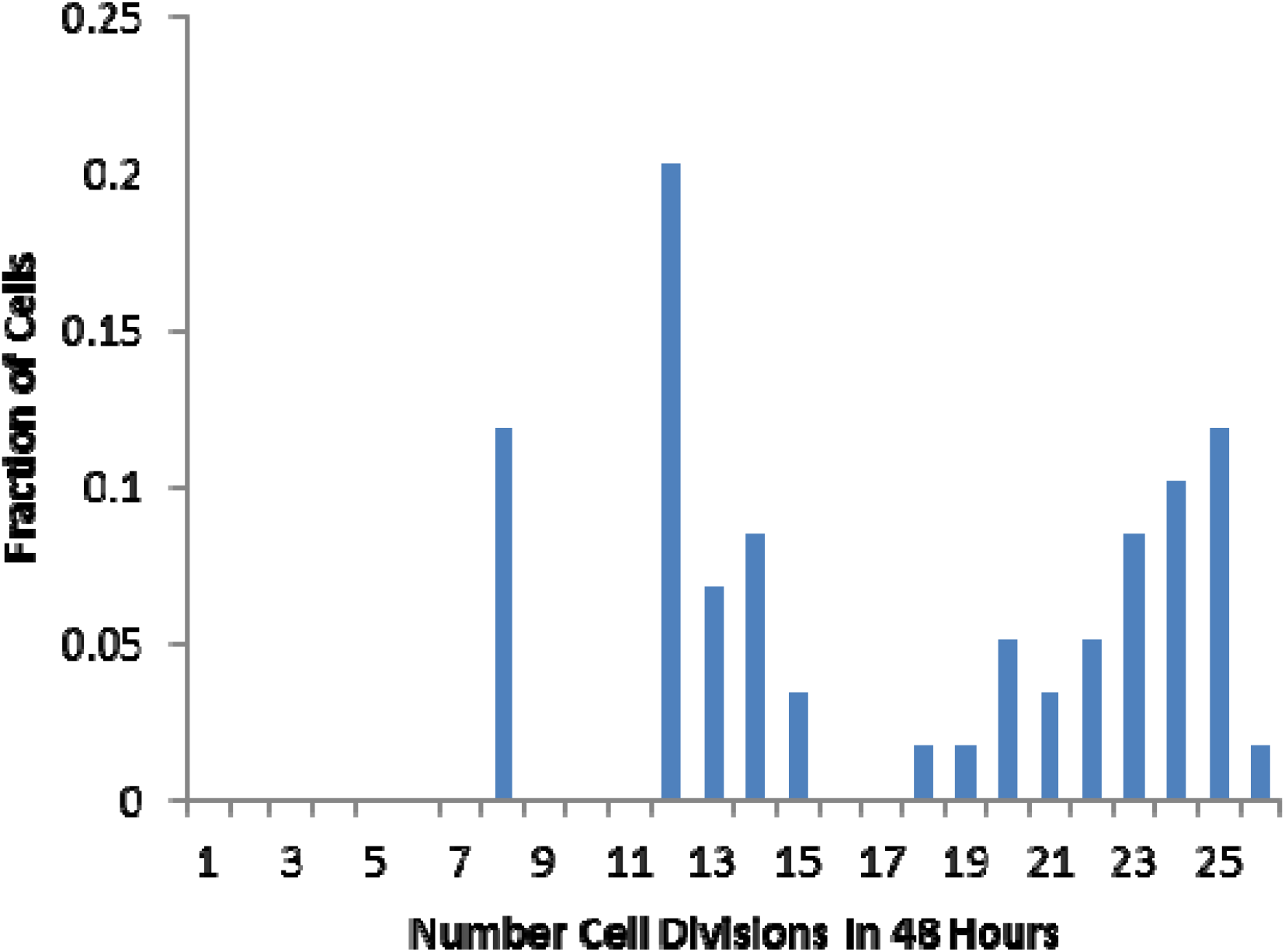
Formation of micro and macrocolonies from cells expressing tub1-828. The number of cell divisions in 48 hours was calculated from the diameter of colonies formed from stationary state cells that had been plated on YPD/tetracycline.

The formation of micro and microcolonies indicates that cell division can occur despite tub1-828’s toxicity. Cells taken from a colony on a refrigerated plate, or from liquid cultures in early and late log phase, or at a stationary-state for 48-360 hours, all displayed this behavior; fewest macrocolonies were formed with log phase cells. Most cells within macrocolonies were not viable (i.e., failed to regrow on YPD minus tetracycline); the fraction of dead cells varied widely and in many cases was c. 90%. Sequencing of genomic DNA from viable cells in macrocolonies revealed that tub1-828 had not undergone reversion to TUB1. Proof that the tub1-828 gene sequence had been retained also came from finding that re-plated viable cells from a macrocolony formed both microcolonies and macrocolonies (Fig. S1, Table S1).

The growth of tub1-828 expressing cells in liquid YPD-tetracycline media was also unusual: growth was greater in cultures started with a large inoculum (Fig. 3A,B, Table 1). This behavior was observed in approximately 25 experiments and was not observed with cells grown in the absence of tetracycline. Cells die more rapidly with a small inoculum because the rate of chromosome loss/cell division was 18-fold greater in a culture with a small inoculum (5000 cells), compared to a large inoculum (350000 cells, Table 1). As noted above, loss of a chromosome other than Chromosome III is lethal in these haploid cells.

**Table 1.**
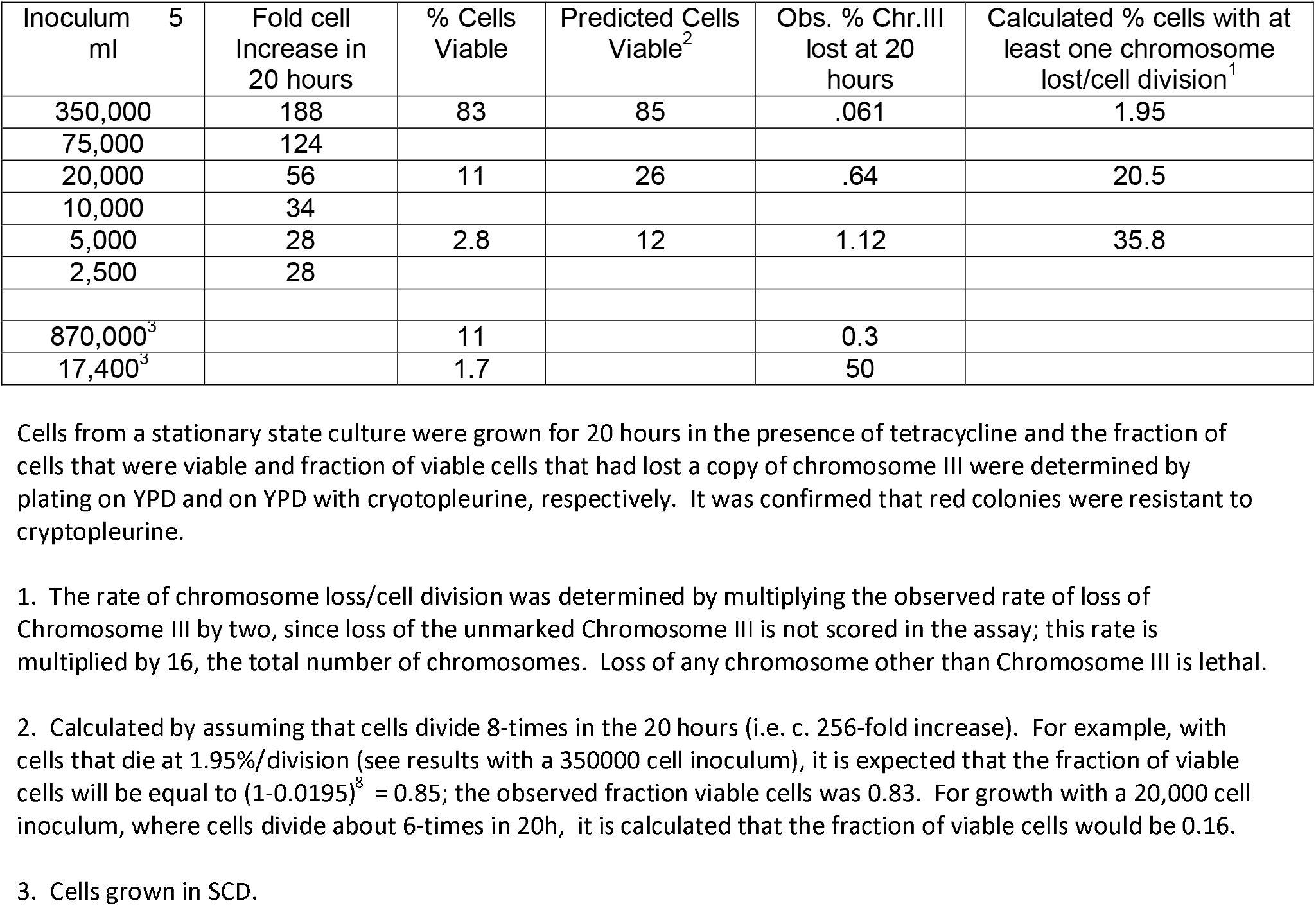
Effect of Cell Density During Expression of tub1-828 on the Loss of Chromosomes

**Fig. 3A.**
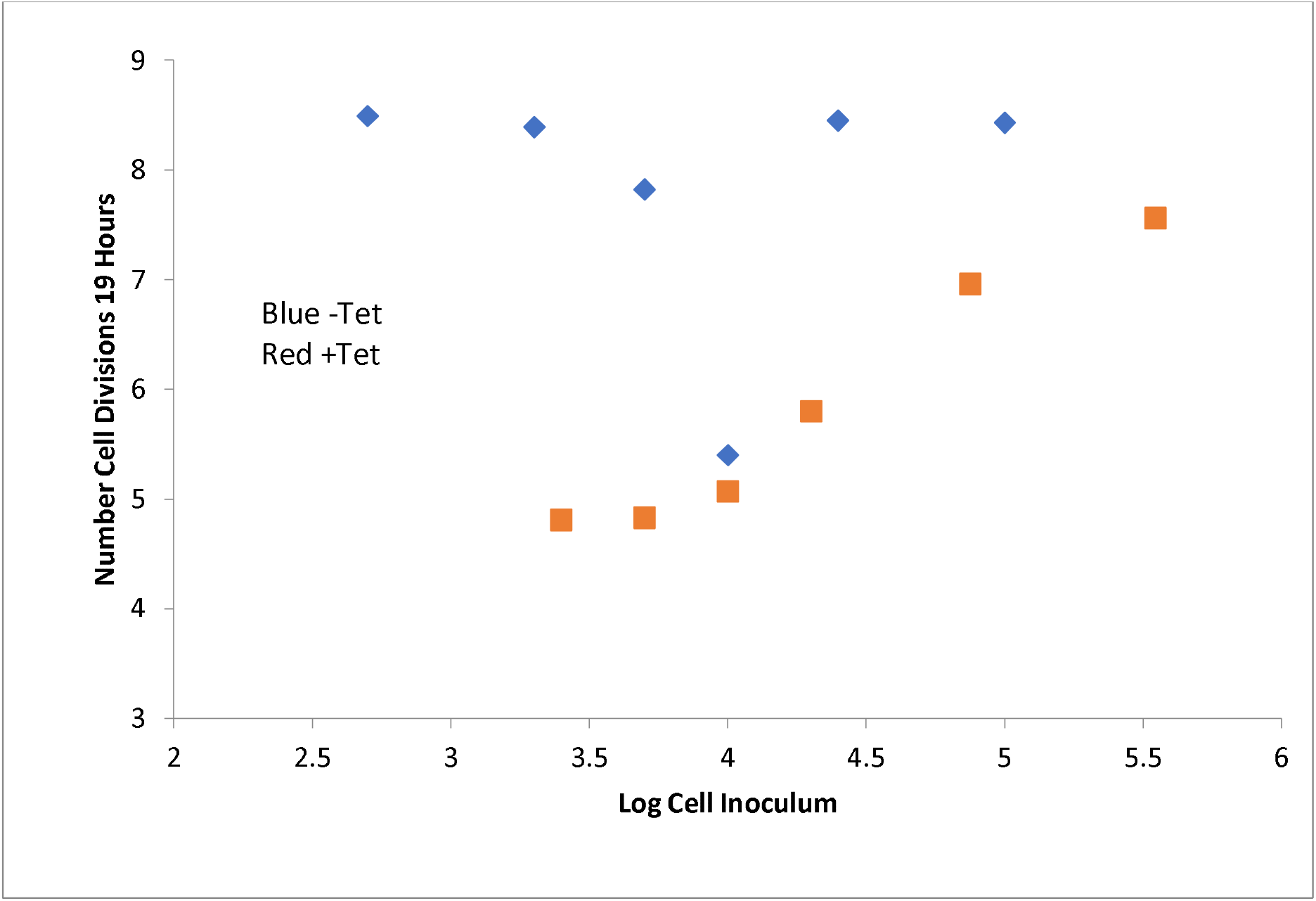
Effect of inoculum size on cell growth in the presence of tetracycline. Cells grown for 72-360 hours were regrown for 19 hours in the presence and absence of tetracycline; the number of apparent cells division was determined with a hemocytometer. Growth was limited by media exhaustion in all of the reactions without tetracycline.

**Fig. 3B.**
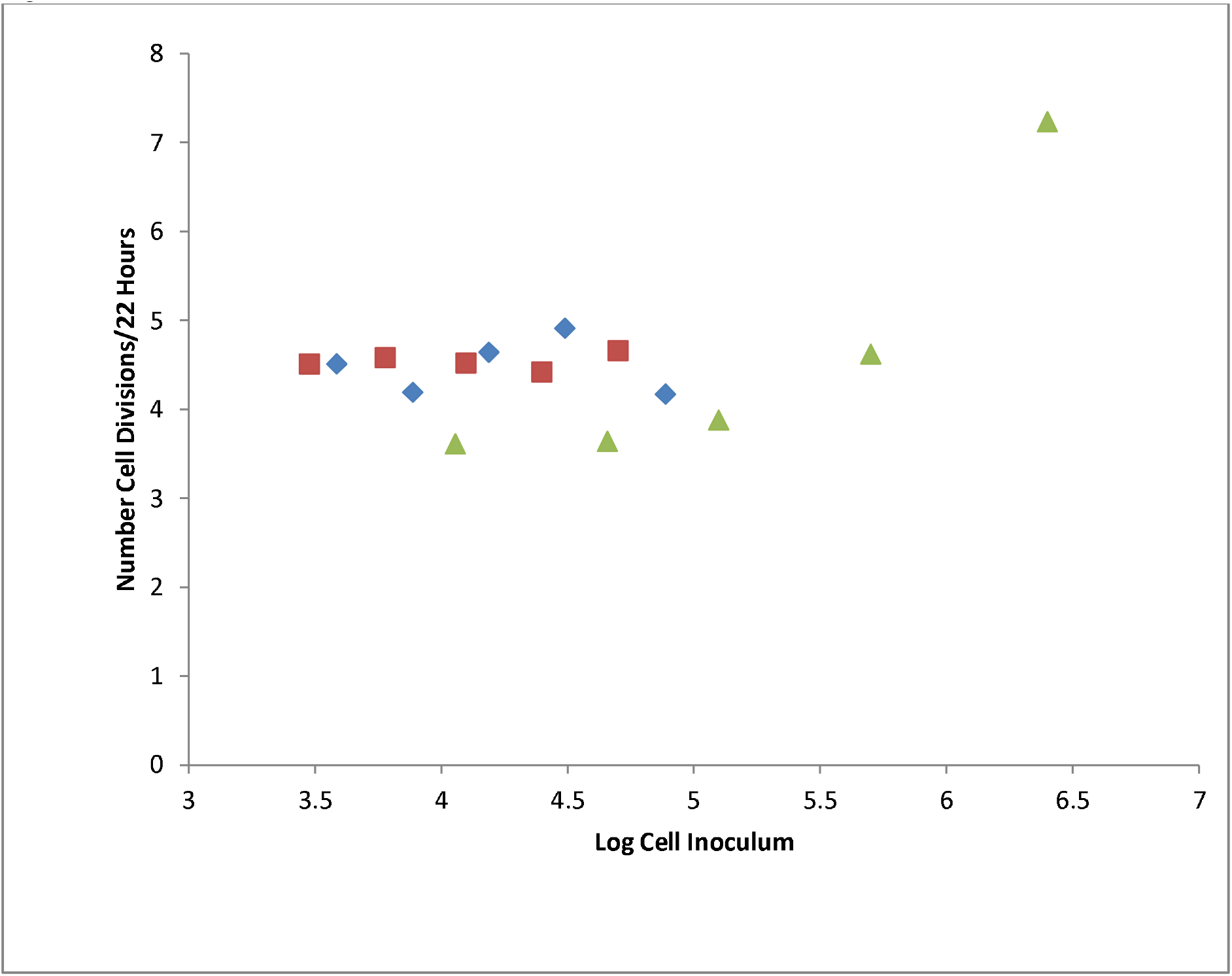
Effect of inoculum size on growth of log phase cells in the presence of tetracycline. Results from three experiments are shown. In the experiment described with green symbols the supernumerary copy of Chromosome III was lost at a rate of 1/343 cells/division and 1/910 cells/division in the reactions at the highest and next-to-highest cell inoculum.

A few studies were done in SCD-tetracycline media where tub1-828 toxicity was much greater: after 20 hours of growth from a 17400 cell inoculum of stationary state cells only 1.7% of cells were viable and 50% of cells had lost the supernumerary copy of chromosome III; with a 870,000 cell inoculum 11% of cells were viable and 0.3% of cells had lost chromosome III (Table 1). It is proposed that slower growth in nutrient-poor SCD prevents rapid accumulation of TUB1 to dilute the toxic tubulin.

Growth from an inoculum of log phase cells was also dependent on the inoculum size (Fig. 3B); however, the number of cells/5 ml to observe stimulation was >125000, compared to 10000 with stationary-state cells. The dependence of growth on cell density suggested that cells secrete a factor that mitigates tub1-828 toxicity, presumably by promoting growth-associated TUB1 synthesis (Fig. 1). The different behavior of log-phase cells may result because they secrete less growth-promoter, or are less sensitive to this factor.

Growth promotion by cells that don’t express mutant tubulin: To obtain additional evidence that cells secrete a growth factor that promotes resistance to tub1-828, we compared growth from a small inoculum of stationary-state cells that express tub1-828 (3440 cells/5 ml) in the absence and presence of 2.3 million stationary-state cells that were otherwise identical, except for lacking the URA-tub1-828 genomic insert. The supplemented cells could not form colonies on agar lacking uracil, so enhanced resistance to tub1-828 expression could be determined from an increased number of URA^+^ colonies with aliquots from cultures with the supplemented cells (Table 2). With cells grown for 6 hours, which is sufficient for three cell doublings, cells at high density underwent approximately one additional cell doubling. This modest effect was anticipated, since stimulation is only expected for the first few cell divisions when cells are induced to grow in fresh media.

**Table 2.** Effect of Heterologous Cells on the Initial Rate of Growth of Cells Expressing tub1-828.

| Reaction | MCY046 Cells | ur Cells | Colonies |
| --- | --- | --- | --- |
| 1 | 3440 | 2360000 | 670 ± 100 |
| 2 | 3440 | 0 | 346 ± 120 |

A small inoculum (3440 cells) of stationary-state MCY046 cells (URA, tet-tub1-828) was grown in 5 ml YPD/tet media with (Reaction 1) and without (Reaction 2) addition of 2360000 cells that were otherwise identical, except that they were ura-minus and did not express tub1-828. Cells were isolated by centrifugation. To be sure that there was equal pelleting of the low number of MCY046 cells in the two reactions 2360000 ura-cells were added to Reaction 2 immediately before cells were isolated by centrifugation. Pelleted cells were washed three times (to remove YPD) before plating on media without uracil. The colonies counted did not include those from ura-minus cells since no colonies were formed when 2360000 ura-minus cells were plated alone.

### Effect of cell density and ATP on the G1-G2 transition

FACS analysis *(Haas and Reed, 2002*) showed the rate of the transition from stationary phase to growth was more rapid with cells grown at high cell density (Fig. 4, Left Panel), and with cells grown with ATP (Right Panel). In this experiment 1.06 E6 stationary-state cells were transferred to 5 ml (high density growth), and 2.12 E5 cells were transferred to 250 ml fresh media (low density growth). Whereas a significant fraction of cells grown at high density duplicated their DNA in 4.5 hours, cells grown at low density were either in G1 or in the process of duplicating their chromosomes, and only a small fraction of cells were in G2. The enhanced growth produced by high cell density was reproduced with cells grown at low density in the presence of ATP (Fig. 4, Right Panel, C and D).

**Fig. 4.**
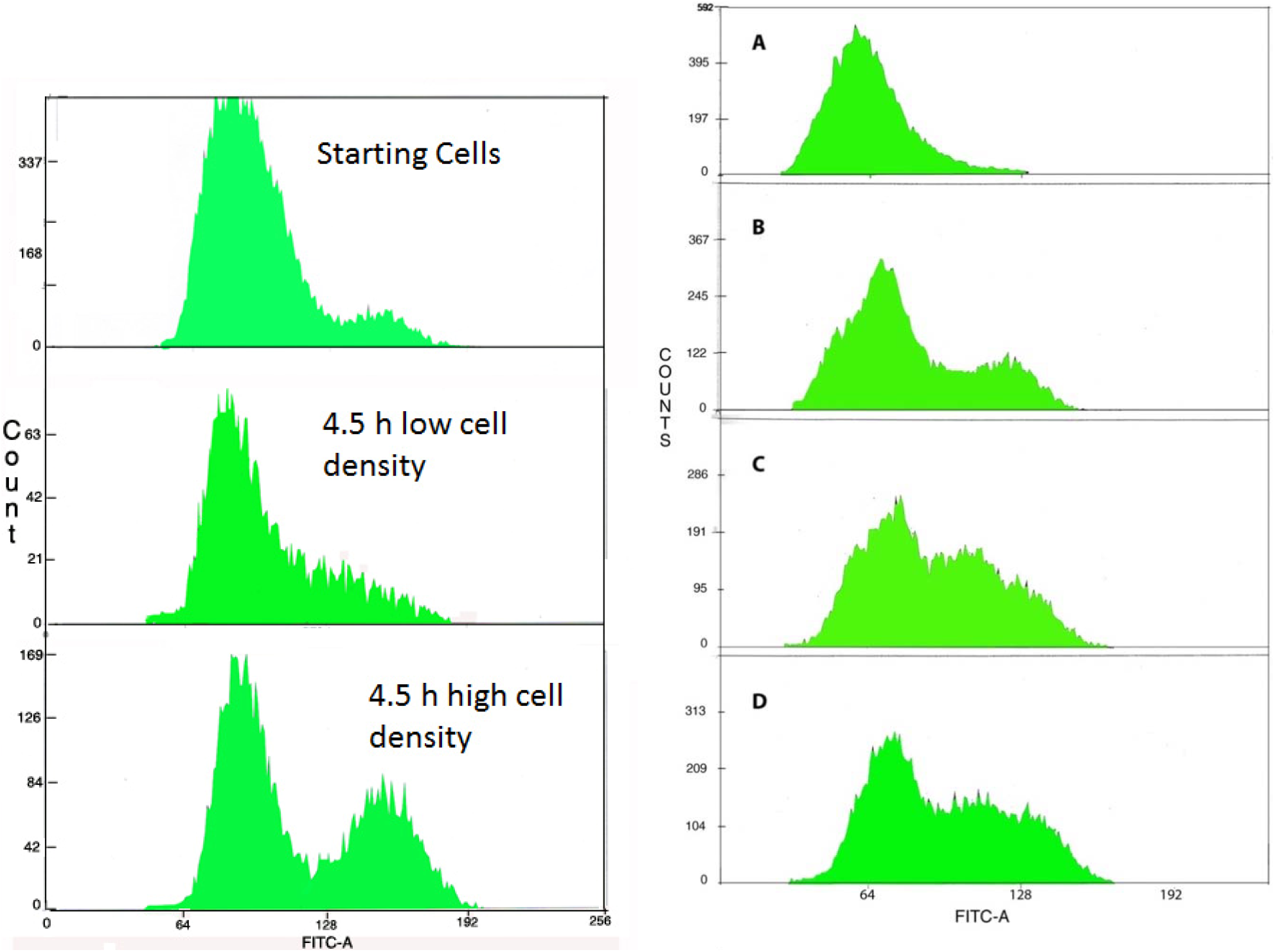
Left DNA profile of stationary-state cells grown at low and at high cell density. The left-hand and right-hand peaks from the cell sorting correspond to cells in G1 and G2; cells that fall between these peaks are in the process of chromosome replication. **Fig. 4 Right Effect of nucleotides on the G1/G2 transition**. Stationary state cells (A: zero-time) were grown in the absence of nucleotide for 5 hours at low density (B) and at high density (C). Growth at low density in the presence of 100 uM ATP (D) was similar to that at high density in the absence of nucleotide (C). Results with CTP (curve not shown) were identical to that for ATP.

#### Nucleotide Effects On Growth

Nucleotides also stimulated growth on tetracycline/agar (Figs. 5, 6, S2, Table S2): For example, after 44 hours, colonies formed from stationary state cells were larger with 100 uM ATP, with the increase corresponding to cells undergoing 12.38, rather than 9.52 cell divisions (Table S2). Both ATP and CTP increased growth, with measurable stimulation with 1 uM ATP (Table S2). ATP was without effect on cells grown without expression of the mutant protein.

**Fig. 5A.**
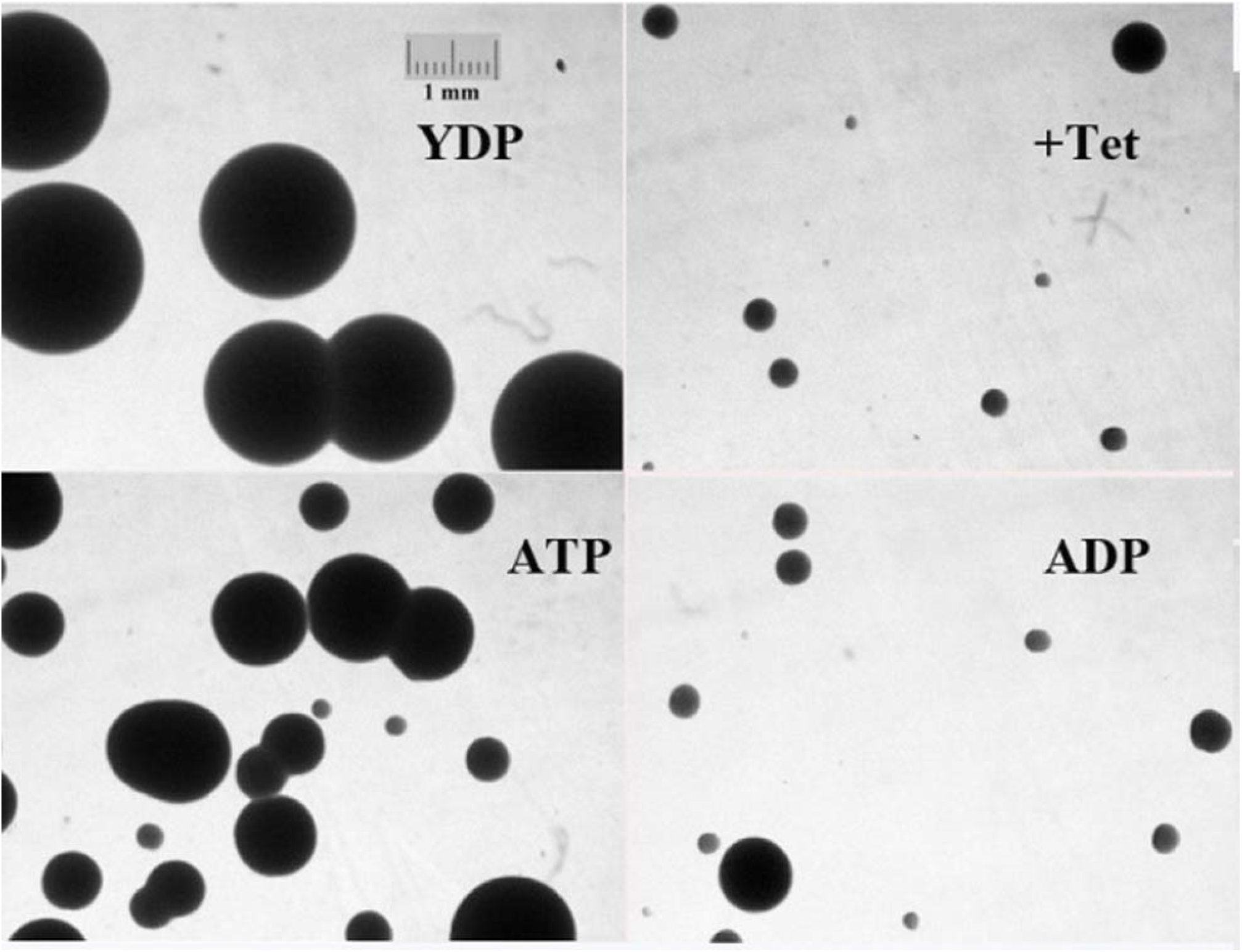
Growth of cells on YPD with and without tetracycline and 100 uM ATP and ADP. The behavior of cells with CTP and GTP and tetracycline was similar to that with ATP.

**Fig. 5B.**
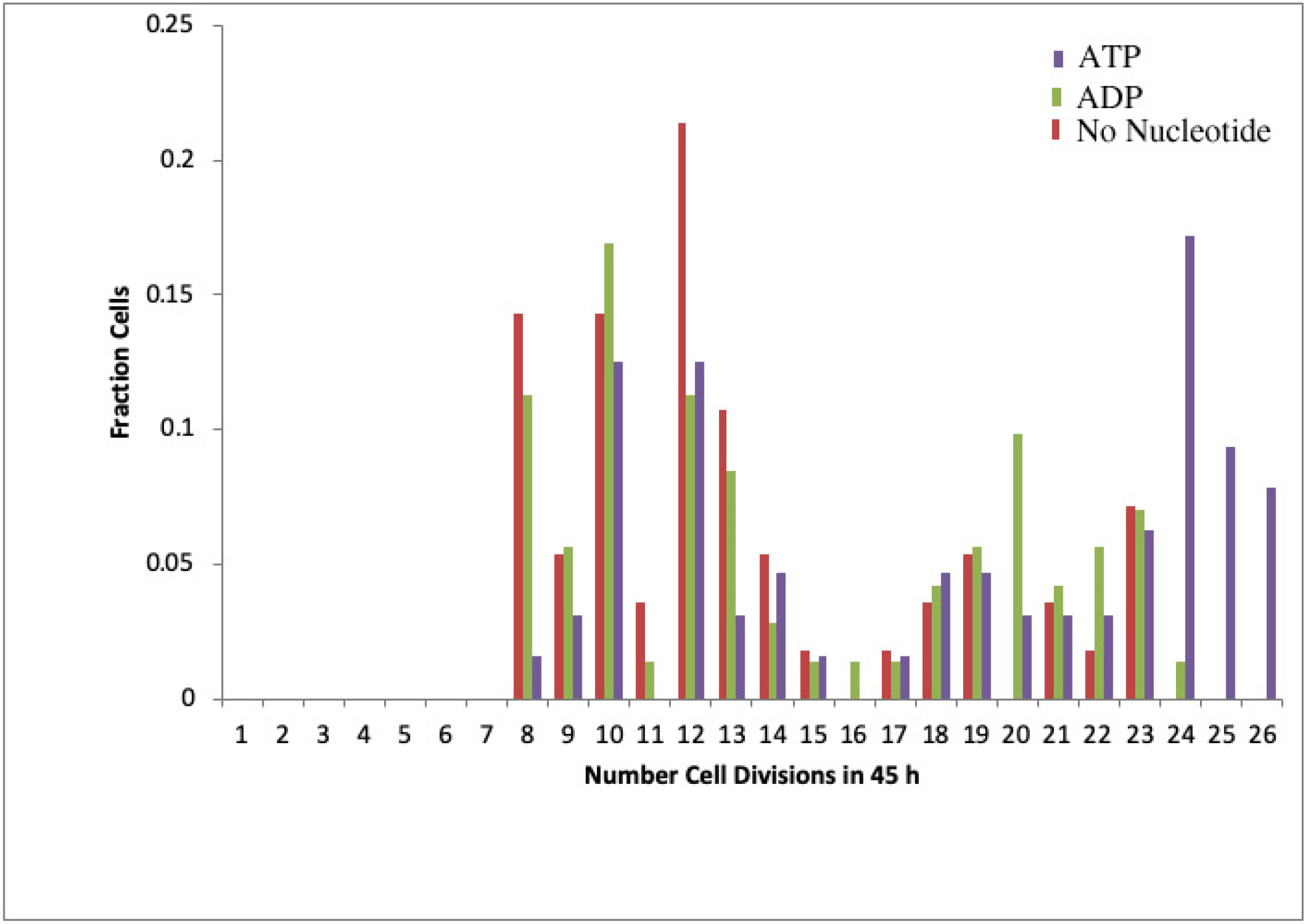
Effect of ATP and ADP on growth of cells expressing tub1-828.

**Fig. 5C.**
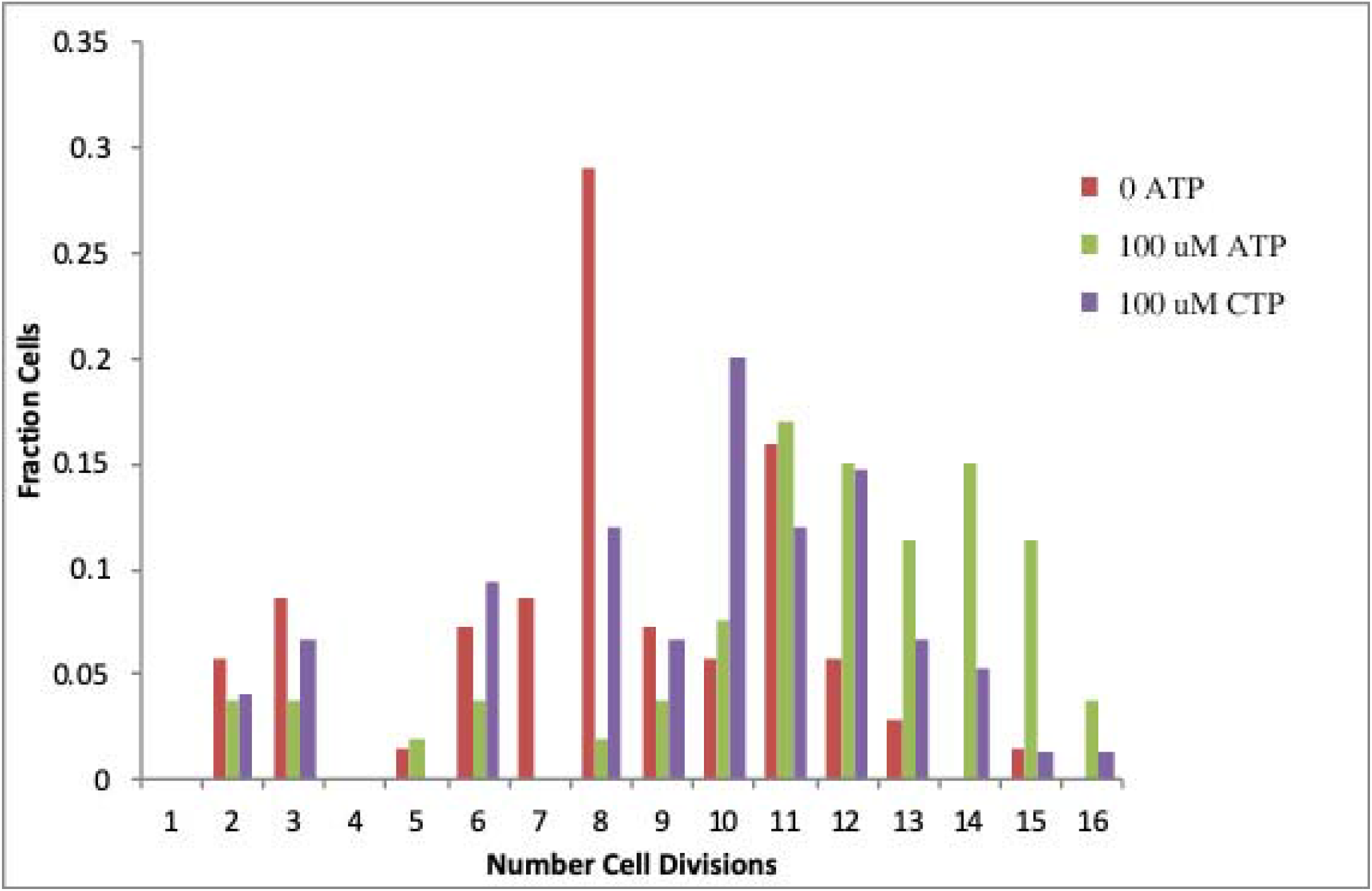
Effect of ATP and CTP on Growth of Cells Expressing tub1-828.

**Fig. 6.**
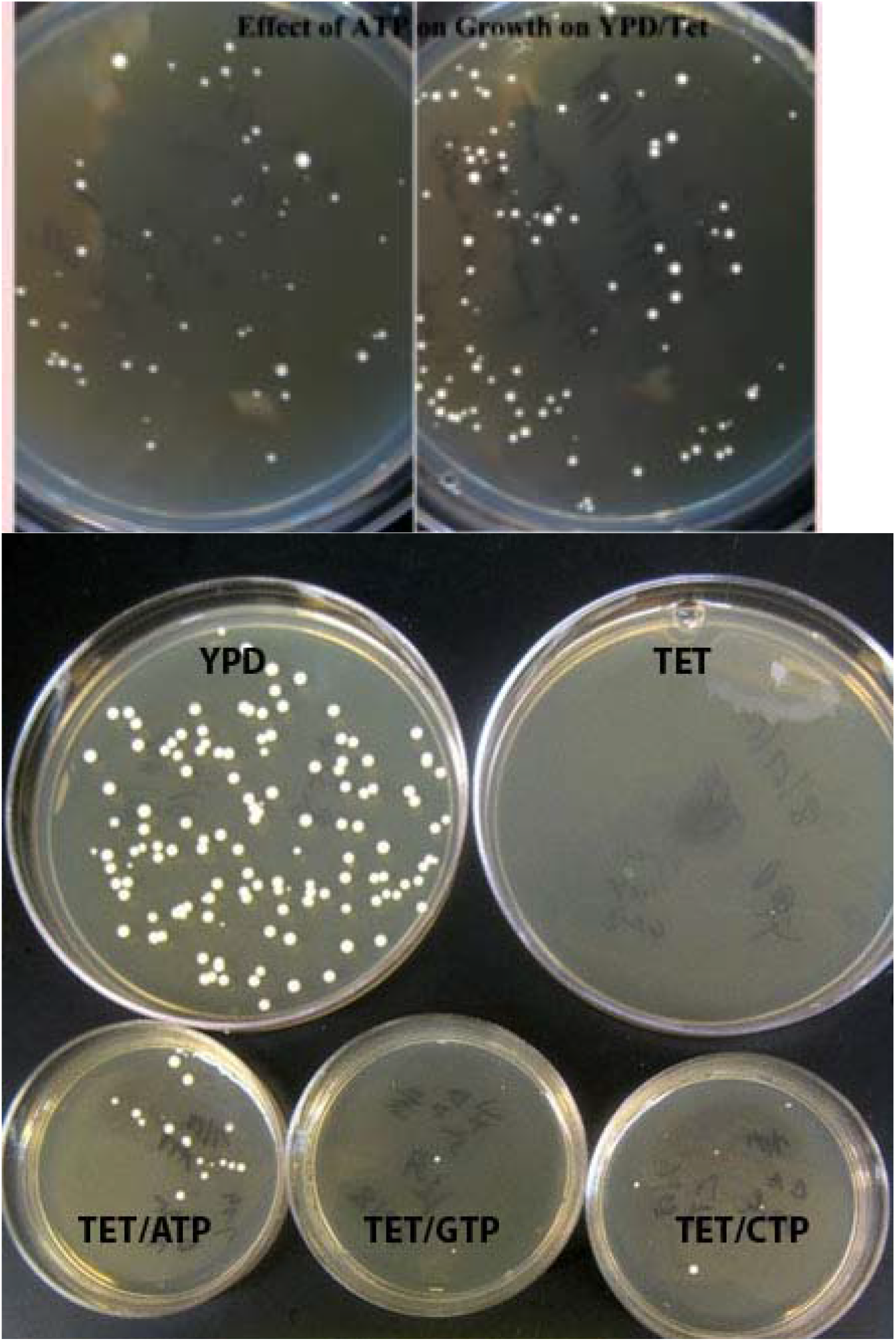
Effect of ATP on growth of cells expressing tub1-828. In this typical experiment 1129 cells that formed colonies on YPD generated 68 macrocolonies (>300 um diameter) on a YPD/Tet plate without ATP (left) and 159 with ATP (right). An additional experiment with ATP, GTP and CTP is below.

As noted above, the behavior of cells from log phase and stationary state cultures in liquid media was different, and this difference was also seen with cells plated on agar. There was only a minimal increase in formation of macrocolonies with cells plated on YPD-tetracycline, with 0.75% of cells forming macrocolonies in the absence of nucleotide and with ATP, and 1.5% with CTP. These nucleotides significantly enhance formation of macrocolonies from stationary-state cells (Figs. 5, 6, S2-3, Table S2).

#### Secretion of ATP

We confirmed earlier results (Boyum and Guidotti, 1997; Esther et al., 2008) showing that cells secrete ATP (Fig. 7). A relatively high concentration of cells generated 20 nM ATP in an hour, corresponding to a rate of about 250 molecules/s/cell; this is similar to that for α-factor (700/s/cell, Goncalves-Sa and Murray, 2011).

**Fig. 7.**
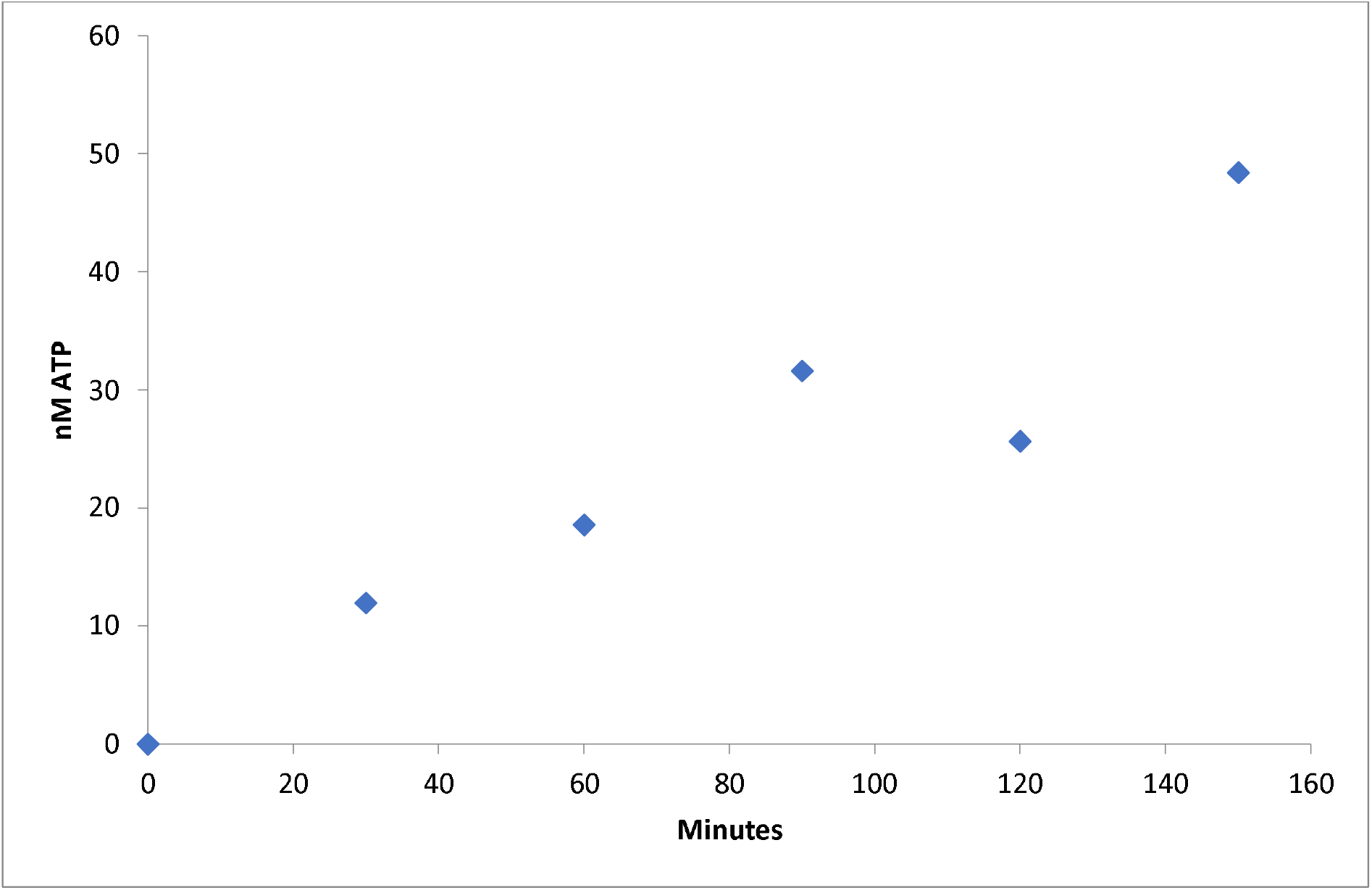
Yeast ATP Secretion. The supernatant from centrifugation of cells (c. 3.7 × 10^7^ /ml in TK buffer (2)) was assayed for ATP as described (2). A comparable rate was seen in YPD in the presence of tetracycline. The rate is approximately 250 molecules/cell/second.

#### Role of Nucleoside Diphosphate Kinase

To determine whether growth stimulation by CTP and GTP (Figs. 5C, 6, S2) resulted from ATP generated by the membrane-associated nucleoside diphosphate kinase’s conversion of ADP to ATP (Zhang et al, 1995), ynk1 cells were analyzed. The results were surprising. First, only about 2% of ynk cells formed microcolonies (Fig. 8), whereas about 50% of YNK cells formed microcolonies, with 14-or fewer divisions, Fig 2B). The predominance of macrocolonies in cells lacking NDP kinase might indicate that this enzyme-signal-transducer normally represses early TUB1 synthesis, so that the null mutant allows early TUB1 synthesis, to repress the effect of tub1-828. The altered behavior of ynk1 cells indicates that NDP kinase regulates cell growth.

**Fig. 8.**
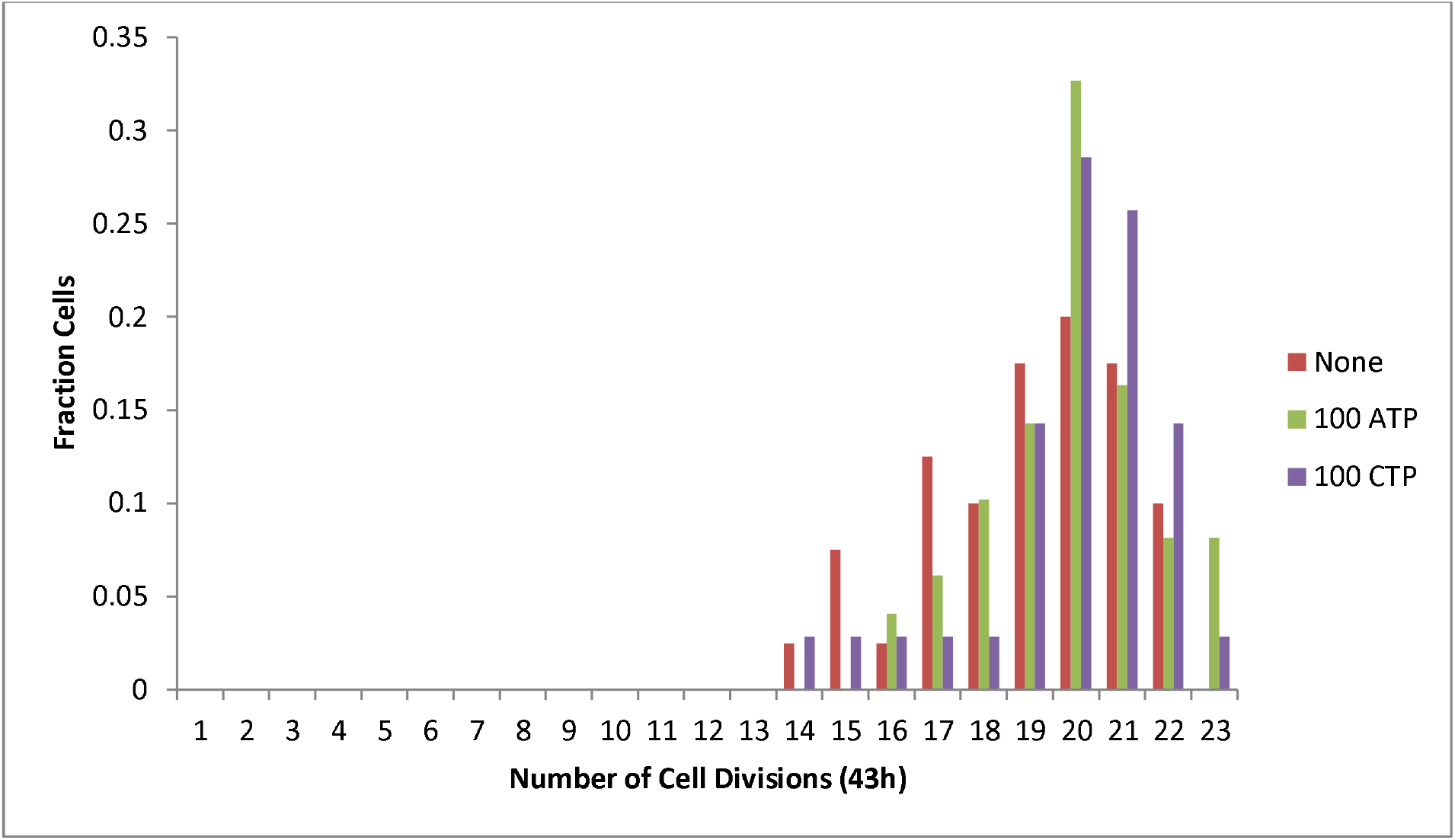
Growth of ynk1 cells in the presence and absence of nucleotides. Cells from a saturated culture divided 19.19, 19.91 and 19.77 times on plates containing without nucleotide and with ATP and with CTP, respectively.

Nucleotides were without effect on growth of an inoculum from ynk1 cells that had been grown to saturation (Figs. 8, S4), suggesting that NDP kinase is part of the NTP signaling path that promotes the transition to cell growth and TUB1 synthesis described above

Although ynk1 cells did not respond to added ATP or CTP in a plating assay, there was greater growth with a larger inoculum for growth in liquid media (Fig. 9). However, as described below, assay of cell growth as a function of the inoculum size is sufficiently sensitive to detect stimulation of 1 out of 1000 cells, so that a stimulatory effect of a secreted growth promoter can be detected by measuring the number of cell divisions before cell death. Such a minuscule stimulatory effect by nucleotides cannot be detected from counting the number or size of colonies on tetracycline plates.

**Fig. 9.**
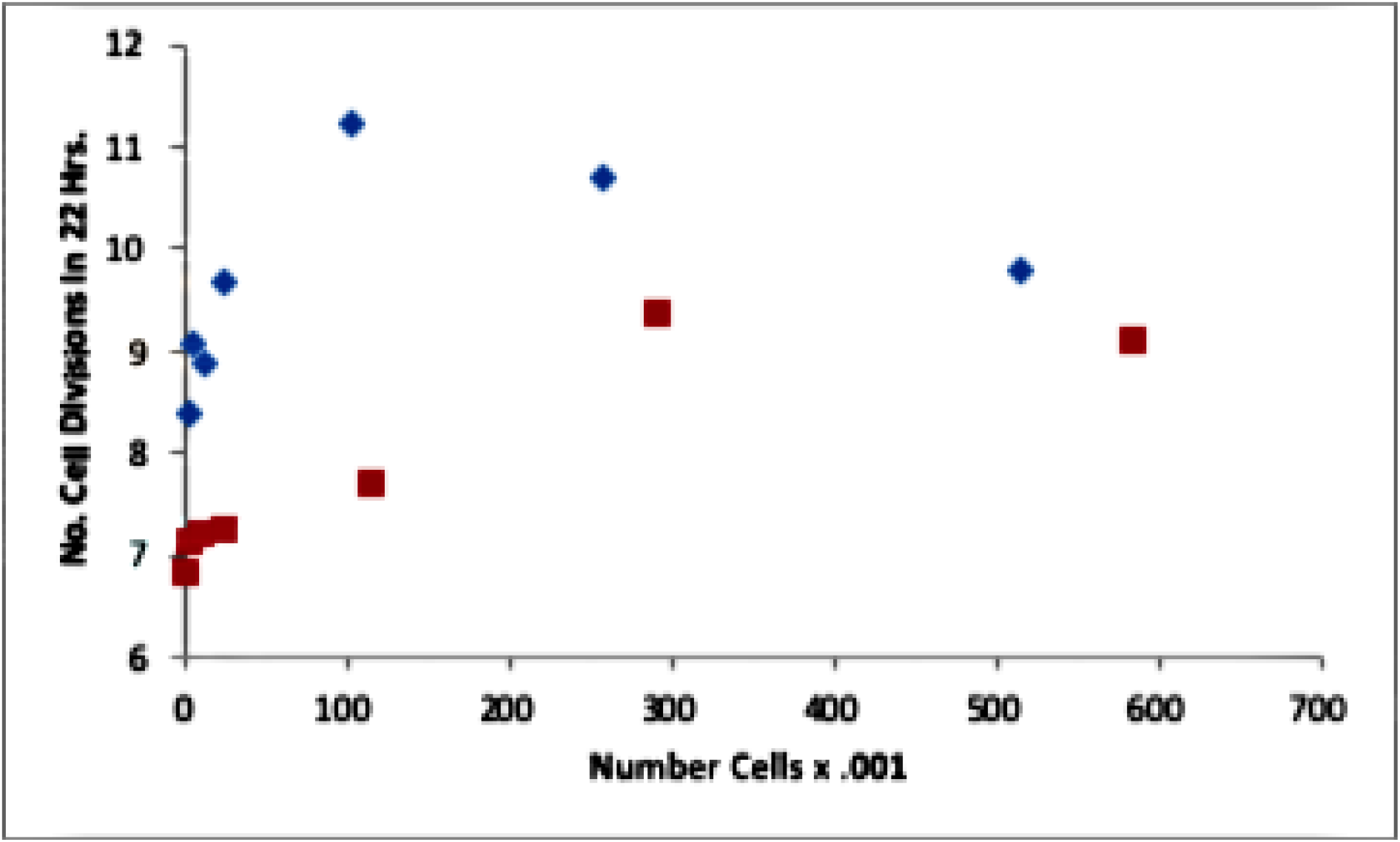
Effect of inoculum size on growth of ynk1 cells (MCY048). Results from two experiments are given.

## Discussion

### Evidence for secretion of a growth substance

It was no surprise to find that induced mutant tub1-828 disrupts the mitotic spindle to allow inaccurate chromosome distribution and cell death (Table 1). What was surprising was that this effect was dramatically reduced in cells grown at high density (Table 1). This presumably results because high cell density produces a sufficient concentration of a secreted signal molecule that stimulates growth and synthesis of wild-type TUB1; TUB1 expression can reduce tub1-828 toxicity (Fig.1).

Additional evidence for secretion of a signal molecule came from the sparing effect of a high density of cells that do not produce tub1-828, on the survival of a low density of tub1-828-expressing cells (Table 2). Similarly, the G1-G2 transition was accelerated in cells at high density (Fig 4); the transition is expected to be accompanied by TUB1 synthesis.

#### A proposed mechanism for resistance to tub1-828 toxicity

Secreted ATP (Boyum and Guidotti, 1997, Esther et al., 2008; Fig 7) is suggested to be the signal molecule for enhanced cell growth and the accompanying TUB1 synthesis. Support for this thesis comes from finding that added ATP reduced tub1-828 toxicity (Fig. 5-6) and accelerated the G1-G2 transition (Fig. 4).

It is postulated that ATP induces rapid growth and earlier than normal TUB1 synthesis, which ordinarily peaks at the entry to mitosis (Spellman et al., 2000), to dilute quickly the tetracycline-regulated toxic protein (Belli et al., 1998). Cells that can generate and sustain a high TUB1/tub1-828 ratio will have greater survival (Fig.1) in both liquid culture (Fig 6A) and on agar (Fig. 2, 10). An increase in the TUB1/tub1-828 ratio can come about if ATP alters the subunit-induced suppression of tubulin mRNA degradation that coordinates TUB1 and TUB2 synthesis (Cleveland et al., 1981), or if ATP induces earlier than normal initiation of TUB1 synthesis.

**Fig. 10.**
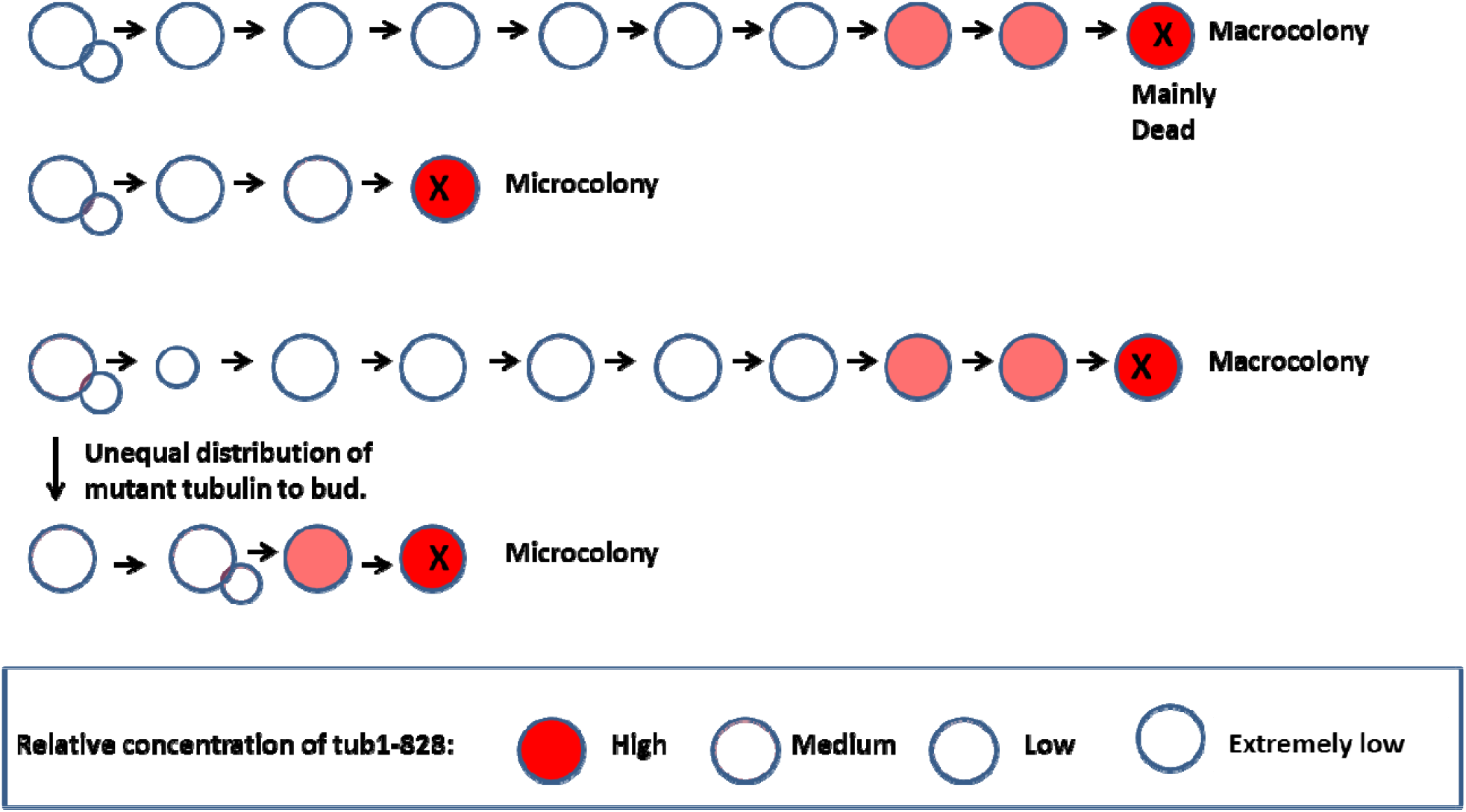
Path for formation of macro and microcolonies during expression of tub1-828. Macrocolonies form if growth (and TUB1 expression) start immediately so that tub1-828 is diluted during the early phase for cell growth.

The wide variation in cell survival (both micro and macro colonies are formed) is presumed to result from cells being randomly distributed in the cell cycle when tub1-828 was induced. The large (30%, Spellman et al., 1998) standard error for TUB1 expression suggests that there are significant variations in TUB1 synthesis during the cell cycle. Cells that are actively synthesizing TUB1 will tend to resist tub1-828 toxicity.

Evidence that cell death is random and precipitous, occurring in the last few divisions, was the fact that macrocolonies were always perfectly spherical (tens of thousands of colonies were measured); if death were progressive, colonies would have been scalloped and misshapen. Also, if death were progressive it would not be possible in 24 hours to generate million-cell-macrocolonies in which 50-90% of cells were dead; persistent cell division is require to generate a million cells. In summary, both the growth of cells that go on to form macrocolonies, and the precipitous death of cells in a microcolony appear to be random events.

Microcolonies were different, although the majority were spherical, about 25% had a shape that suggested that death occurred in an early point, so that a large portion of the sphere was missing. A possible basis for early and precipitous cell death from tub1-828 is discussed next.

#### Cooperativity for tub1-828 toxicity

Our earlier study of the concentration-dependence of the effect of tubulin subunits containing non-hydrolyzable GMPCPP (Caplow and Shanks, 1996) may explain how tubulin with unhydrolyzed GTP (tub1-828 subunits) effects microtubules.

Microtubules are capped and transiently stabilize by tubulin-GTP subunits (Desai and Mitchison, 1997, Roostalu et al., 2020). Although the size of the stabilizing GTP-cap is not known, in vitro studies of microtubules with tubulin subunits containing a nonhydrolyzable GTP analogue (GMPCPP) demonstrated that stabilization requires 13 or 14 contiguous tub1-828 subunits at a microtubule end (Drechsel and Kirschner, 1994; Caplow and Shanks, 1996). This means that microtubules stabilization by tub1-828 subunits, and the resultant toxicity will be proportional to the 13^th^ or 14^th^ power of the mole fraction of tub1-828 subunits in the cell’s pool of alpha-tubulin. Therefore, a slight decrease in TUB1 expression, or a small stochastic increase in tub1-828 expression in individual cells (Raser and O’Shea, 2005; Bar-Even et al., 2006) will, after a variable number of cell divisions, result in loss of one of the cell’s 16 chromosomes, and cell death.

Our finding increased cell survival (Fig. 6A) at cell densities that form low concentrations of ATP (Fig. 7) may be accounted for by the just-described mechanism. With microtubule stabilization having a 13^th^ or 14^th^ power dependence on the mole fraction of tub-818 subunits, even minimal stimulation of cell growth is likely to generate sufficient TUB1-GTP subunits to disrupt the purity of a 14-subunit tub1-828-GTP cap. Thus, the dependence of cell survival on ATP concentration does not reflect a Kd for ATP binding, but rather a measure of its stimulation of TUB1 synthesis to disrupt the effect of a tub1-828 cap.

#### Correlation of ATP Secretion with Cell Growth

It is suggested that secreted ATP suppresses toxicity of tub1-828 by a paracrine mechanism for cells grown in liquid media and by an autocrine mechanism for cells grown on agar.

Is sufficient ATP generated in liquid media to stimulate growth: Although yeast generate 1 uM ATP in 40 hours (Peters et al., 2016), a lesser amount is secreted during the first 30-60 minutes of growth, when the postulated stimulation of TUB1 synthesis is required to offset tub1-828 toxicity. However, as described next, there is a huge multiplier effect that allows detection of minimal growth stimulation by low ATP concentrations. For example, if the ATP concentration is only sufficient to stimulate 1 of 1000 cells, and the unstimulated 999 die after only 10 division (i.e., form what would be microcolonies in an agar assay), a total of 1.02 million cells result; however, if the 1 cell that responds to ATP is able to divide 20-times (i.e. form what would be a macrocolony in an agar assay), 1.05 million cells are generated. Thus, ATP stimulation of 1 cell in 1000 increases the final cell yield from 1.02 million to 2.07 million; i.e., there would appear to be one extra cell division. The result from this analysis reasonably agrees with our observation: with 100,000 cells/5 ml (Fig. 3A) the calculated (Fig. 7) ATP concentration after one hour would be about 0.03 nM. Since 1 uM ATP has a detectable effect using the enormously less sensitive agar-plating assay, it is likely that 0.03 nM ATP would produce the results in Fig. 3A. Finally, our finding growth stimulation by extracellular ATP concentrations 1000-fold lower than the 1.3 mM ATP concentration in cells (1.3 mM, Ytting et al., 2012) indicates that extracellular ATP signaling occurs before ATP enters the cell.

For single cells plated on agar, the same question arises, whether sufficient ATP is produced to stimulate growth. However, the situation is different because of reduced diffusion by both agar, and an unstirred layer surrounding the cell membrane (USL; Gabba et al. 2020). Yeast and other cells in stirred liquid media have an USL that provides diffusive resistance to uptake of added permeating substances, and loss, by diffusion, of secreted substances (Gabba et al., 2020). The USL for a cells on agar is expected to be especially stable since media stirring, which reduces the USL (Pedley, 1983), is absent.

A rough estimate of the USL steady-state ATP concentration comes from the ATP secretion rate and the volume of the USL. It is suggested that the volume corresponds to that of the approximately 100 um cell wall (Dupres et al., 2010). Assuming a 4.8 um diameter for the volume enclosed by the plasma membrane (5.79 E-14 L), and a 5.0 um diameter for the volume enclosed by the cell wall (6.54 E-14), the volume within the cell wall is 7.5 E-15 L. The ATP concentration contained in the USL is expected to be at a steady state both before and after a cell is plated. Assuming this steady-state corresponds to the continuous secretion of ATP at a rate of 250/s, and an equal loss of ATP by diffusion away from the USL. The estimated steady-state concentration equal 250/7.5 E-15 = 0.05 uM, which is sufficient to activate the small fraction of cells to form macrocolonies on ATP-free agar.

#### Role of nucleoside diphosphate kinase in growth stimulation

Cells made null for nucleoside diphosphate kinase (ynk1) have an altered growth response to tub1-828 and an altered response to nucleotides. Induction of tub1-828 from stationary state cells resulted in rapid growth, without formation of microcolonies. This can be taken to indicate that NDP kinase negatively regulates early TUB1 synthesis and growth so that in a null mutant sufficient TUB1 is synthesized to offset tub1-828 toxicity. NTP stimulation of TUB1 synthesis is obviously not required for growth. The very different results with quiescent ynk1 cells, which are sensitive to induced tub1-828, allowed test of an hypothesis that NDP kinase is required for NTP stimulation of growth: results supported this hypothesis (Fig. 9). This conclusion is not controverted by results showing a dependence of growth if ynk1 cells on the inoculum size (Fig. 10). As noted above, assay for growth stimulation in liquid media is enormously more sensitive to growth stimulation, compared to the cell plating assay used in Fig 9: enhanced growth by 1 cell out of 1000 can be detected. Our finding that NDP kinase has a regulatory role for cell-cell communication and growth is in accord with observations that this protein regulates tumor metastasis and drosophila development and sexual development in S. pombe (Izumiya and Yamamoto, 1995; Woolworth et al. 2009).

#### Yeast signaling

Discovery of an extracellular ATP signaling pathway for stimulating yeast growth adds to the S. cerevisiae signaling paths for: quorum sensing and generation of an undifferentiated multicellular species (Chen and Fink, 2006); for the dimorphic transition induced by ammonia (Vopálenská et al., 2010); and for sexual reproduction. Although yeast lack a purinergic receptor, evidence that this does not necessarily preclude an ability to respond to ATP comes from a recent finding uM concetrations of exogenous ATP influence the nitrogen stress response of S pombe (Forte et a, 2019). Also, plants, which like yeast lack purinergic receptors, use ATP to regulate growth, development and the response to stress (Demidchik et al, 2003; Choi et al., 2014).

The signaling pathway for ATP stimulation of the transition from stationary-state to rapid growth is unknown, but may involve activation of the Yck1p and/or the YCK2 membrane-associated casein kinases that mediate the transition from quiescence to rapid growth in starved cells (Zaman et al., 2008; Wand et al., 1996). The ATP-binding domain for these enzymes may require extracellular ATP for activity. Alternatively, the ATP signaling path may involve YNK1. Evidence for this was the absence of an ATP effect on growth on tetracycline plates with cells that are nul for this enzyme-signaling protein (Izumiya and Yamamoto, 1995; Woolworth et al. 2009).

Our studies indicate that yeast decision-making for adapting to a new environment is not entirely autonomous. Information provided by the ATP growth-promoting signal is somehow integrated into the complex regulatory mechanisms that induce the vast changes in protein expression that allows adaptation to new conditions (Gasch et al., 2000; Gelade et al., 2003).

Stimulation by secreted ATP of growth of neighboring cells in the wild may be of utility since upon encounter with nutrients an increase in a colony’s population will serve to diminish food for neighboring rival organisms, and thereby increase its likelihood of survival.

## Acknowledgements

A colleague, Howard Fried provided greatly valued encouragement throughout this work, as well as valuable practical advice. I also profited from Henrik Dohlman’s generosity and helpfulness in informing me that yeast secrete ATP and from Leslie Parise and Nikolay Dokholyan, who generously provided lab space.

## Notes

### Competing Interest Statement

The authors have declared no competing interest.

