## Supplementary material for "ATP Mediates Yeast Cell-Cell Communication": 032222 Supplementary Material

**S1** **Cells from a macrocolony form both micro and macrocolonies when grown for 48 hours on YPD.** Results from cells from two macrocolonies are shown.

Fraction Cells

| No. Cell Divisions | Percent |
| --- | --- |
| 1 | 0 |
| 2 | 0 |
| 3 | 0 |
| 4 | 0 |
| 5 | 0.241379 |
| 6 | 0 |
| 7 | 0 |
| 8 | 0 |
| 9 | 0.310345 |
| 10 | 0.206897 |
| 11 | 0.0689655 |
| 12 | 0.0689655 |
| 13 | 0 |
| 14 | 0 |
| 15 | 0.0344828 |
| 16 | 0 |
| 17 | 0.0344828 |
| 18 | 0.0344828 |

Fraction Cells

| No. Cell Divisions | Percent |
| --- | --- |
| 1 | 0.0645161 |
| 2 | 0.129032 |
| 3 | 0 |
| 4 | 0 |
| 5 | 0 |
| 6 | 0 |
| 7 | 0 |
| 8 | 0 |
| 9 | 0 |
| 10 | 0.0322581 |
| 11 | 0.0322581 |
| 12 | 0 |
| 13 | 0.0322581 |
| 14 | 0 |
| 15 | 0 |
| 16 | 0.0322581 |
| 17 | 0 |
| 18 | 0 |
| 19 | 0 |
| 20 | 0 |
| 21 | 0.0322581 |
| 22 | 0.0322581 |
| 23 | 0.0322581 |
| 24 | 0.0967742 |
| 25 | 0.451613 |
| 26 | 0.0322581 |

**Fig. S2A Nucleotide Effect on Growth**

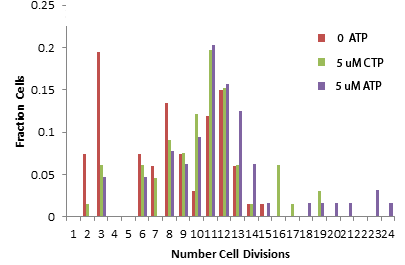

**Fig S2B**

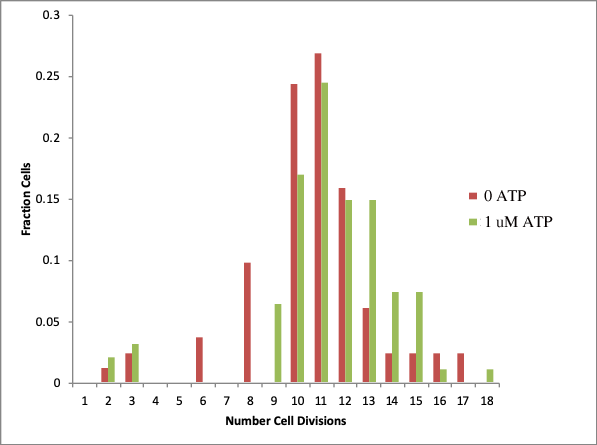

**Fig. S3** **Nucleotide effect on growth of ynk1 cells**. Growth of 616 viable cells (determined by growth in the absence of tetracycline) from a refrigerated colony for 72 hours gave 17 macrocolonies (c. 1 mm diameter) in the absence of nucleotide (A), 26 with 10 uM ATP (B), 25 with 100 uM CTP (C), and 19 with 100 uM ATP (D). The accompanying microcolonies were small (corresponding to 20-50 um, 5-9 cell divisions), without intermediate size colonies.

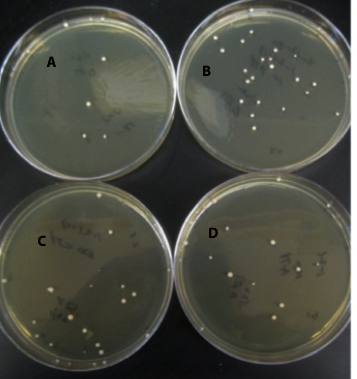

**Table SI**

**Characterization of Cells in Macrocolonies Described in Fig. S1**

| Colony | % Viable | % Cells Yielding Macrocolonies on tet | Cells Losing Chromosome III |
| --- | --- | --- | --- |
| 1 | 34 | 38. | 0/390 |
| 2 | 46 | 0.22 | 5/6900 |

Cells from a macrocolony were re-plated on YPD, YPD-Tet and YPD-cryptopleurine, to determine the percent of cells that were viable, the percent that could form a microcolony, and the percent that had lost the supernumerary Chromosome III, respectively.

**Table S2**

**Effect of Nucleotide Concentration on Colony**

**Growth During tub1-828 Expression**

| Experiment | No. Apparent Cell Divisions | No. Apparent Cell Divisions | No. Apparent Cell Divisions |
| --- | --- | --- | --- |
| 1 | 0 ATP = 9.52 | 100 uM ATP = 12.38 | 100 uM CTP = 10.91 |
| 2****** | 0 ATP= 10.1 | 5 uM ATP = 18.20 | 5 uM CTP = 13.50 |
| 3 | 0 ATP = 11.96 | 1 uM ATP = 12.24 | - |

Stationary-state cells were plated on tetracycline-agar with and without nucleotide and colonies were measured after 44 hours. Nucleotide stimulation is described as an increase in “apparent cell divisions”; i.e., number of divisions in 100 colonies generated from 100 plated cells.

****** A greater number of apparent cell divisions had occurred because colonies were measured after 68 rather than 44 hours.
